# DivIVA is essential in *Deinococcus radiodurans* and its C terminal domain regulates new septum orientation during cell division

**DOI:** 10.1101/2020.04.09.033746

**Authors:** Reema Chaudhary, Swathi Kota, Hari S Misra

## Abstract

FtsZ assembly at mid cell position in rod shaped bacteria is regulated by gradient of MinCDE complex across the poles. In round shaped bacteria, which lack predefined poles and the next plane of cell division is perpendicular to previous plane, the determination of site for FtsZ assembly is intriguing. *Deinococcus radiodurans* a coccus shaped bacterium, is characterized for its extraordinary resistance to DNA damage. Here we report that DivIVA a putative component of Min system in this bacterium (drDivIVA) interacts with cognate cell division and genome segregation proteins. The deletion of full length drDivIVA was found to be indispensable while its C-terminal deletion (Δ*divIVAC*) was dispensable but produced distinguishable phenotypes like slow growth, altered plane for new septum formation and angular septum. Both wild type and mutant showed FtsZ foci formation and their gamma radiation responses were nearly identical. But unlike in wild type, the FtsZ localization in mutant cells was found to be away from orthogonal axis with respect to plane of previous septum. Notably, DivIVA-RFP localizes to membrane during cell division and then perpendicular to previous plane of cell division. *In trans* expression of drDivIVA in Δ*divIVAC* background could restore the wild type pattern of septum formation perpendicular to previous septum. These results suggested that DivIVA is an essential protein in *D. radiodurans* and the C-terminal domain that contributes to its interaction with MinC determines the plane of new septum formation, possibly by controlling MinC oscillation through orthogonal axis in the cells.

## Introduction

Mechanisms underlying cell division is better understood in rod shaped bacteria like *Escherichia coli* and *Bacillus subtilis*. The bacterial cell division is initiated with the polymerization of the tubulin-like protein FtsZ at the mid cell position in rod shaped bacteria (Bi & Lutkenhaus, 1991) and perpendicular to first plane of cell division in cocci (Begg & Donachie, 1998). The proper localization and polymerization of FtsZ are critical events for both initiation of FtsZ ring formation and recruitment of other downstream cell division proteins to form a productive divisome complex (de Boer, 2010; Egan & Vollmer, 2013; Lutkenhaus *et al*., 2012; Rico *et al*., 2013). Many proteins involved in cell division are conserved across the bacterial species except to those involved in spatial and temporal regulation of FtsZ localization (Margolin, 2005; Monahan *et al*., 2014). In *E. coli* and *B. subtilis*, the regulation of FtsZ localization and polymerization at mid cell position is spatially regulated by Min system comprised of MinCDE (de Boer *et al*., 1988, 1989, 1992; Lee & Price, 1993), and temporally by nucleoid occlusion (NOC) mechanisms manifested by SlmA in *E. coli* (Bernhardt & de Boer, 2005) and Noc in *B. subtilis* (Wu & Errington, 2004, 2012; Barak & Wilkinson, 2007; Monahan *et al*., 2014). In Gram-positive bacteria MinE is replaced with DivIVA. Some bacterial species have only one of these systems while many lacks both NOC and Min systems (Margolin W., 2005; Harry *et al*., 2006; Monahan et al., 2014), which indicates towards a possibility of alternate mechanism(s) for both spatial and temporal regulation of FtsZ functions in bacteria. In fact, there are many bacteria that are discovered with alternate mechanisms for the regulation of FtsZ functions. For example, PomZ in *Myxococcus xanthus (*Treuner *et al*., 2013), MipZ in *Caulobacter crescentus* (Thanbichler M. & Shapiro L., 2006), CrgA in *M. smegmatis* (Plocinski P. *et al*., 2011) and Ssd in *M. smegmatis* and *M. tuberculosis* (England K. *et al*., 2011) have been characterized for their regulatory roles in cell division and septum growth. How the previous plane of cell division is memorised and what determines the next plane of division have remained the interesting questions in cocci.

*Deinococcus radiodurans* is a Gram-positive coccus bacterium that exist in tetrad form and divides in alternate orthogonal planes perpendicular to each other (Murray *et al*., 1983). This bacterium is better known for its extraordinary resistance to DNA damage produced by physical and chemical agents including radiations and desiccation (Cox and Battista, 2005, Slade and Radman, 2011; Misra *et al*., 2013). It is noteworthy that it can tolerate nearly 200 double strand breaks and 3000 single strand breaks without loss of cell viability. In *D. radiodurans*, the dividing septum grows in a closing-door mechanism and not as a diaphragm has been reported for other cocci (Touhami *et al*., 2004; Wheeler *et al*., 2011; Floc’h *et al*., 2019). It possesses putative Min system comprised of MinC, MinD, an aberrant MinE and DivIVA whereas the homologues of known NOC systems are not annotated in the genome (White *et al*., 1999). However, a P-loop DNA binding ATPase (ParA2) encoded on chromosome II in *D. radiodurans* has been characterized for NOC-like function (Charaka *et al*., 2013). The DivIVA in *D. radiodurans* (drDivIVA) has two structural domains named N-terminal domain (NTD) and C-terminal domain (CTD). NTD consists of lipid binding motif, which is conserved across Gram-positive bacteria and CTD varies in different bacteria. Recently, it has been shown that the genome segregation proteins interact through NTD while MinC interacts to DivIVA through its CTD that contains middle domain. Apart from this, the other functional significance of full length as well as CTD in the regulation of cell division is not known. Here, we report that DivIVA is an essential protein in *D. radiodurans* and CTD interacts with MinC, which is required for its axial oscillation. Such a process helps in the formation of FtsZ ring perpendicular to MinC oscillation and thus septum growth at 90°C to previous septum. We demonstrated that as the copy number of *drdivIVA* decreases in the cell, the lethality increases and the cells approaching to homogenous replacement of *drdivIVA* with *nptII* cassette get debilitated. The selective removal of C terminal domain (hereafter referred as Δ*divIVAC*) did not affect survival, but growth rate was relatively compromised. Gamma radiation effect was found to be of the same magnitudes as that of wild type cells. These results supported the indispensability of DivIVA in this bacterium and an important role of the CTD of drDivIVA in macromolecular interaction that determines the perpendicularity of alternate plane of cell division is suggested.

## Materials and Methods

### Bacterial strains, plasmids, and growth conditions

*D. radiodurans* R1 (ATCC13939) was a kind gift from Professor J. Ortner, Germany (Schafer M., 2000). It was grown in TGY (1% Tryptone, 0.5% Yeast extract and 0.1% Glucose) medium at 32°C with shaking speed of 160 rpm. *E. coli* NovaBlue was grown in Luria-Bertini (LB) broth (1% Tryptone, 0.5%Yeast extract and 1% sodium chloride) at 37°C with shaking speed of 165 rpm, was used for cloning and maintenance of all the plasmids. *E. coli* cells harbouring different plasmids viz. pNOKOUT (Khairnar *et al*., 2008) and shuttle expression vector pRadgro (Misra *et al*., 2006) and their derivatives were maintained in *E*.*coli* in the presence of kanamycin (25 µg / ml) and ampicillin (100 µg / ml), respectively. *Deinococcus radiodurans* R1 and its mutants were grown in TGY medium in 48 well microtiter plates at 32°C for 42 h. Optical density at 600 nm (OD600) was measured online in the Synergy H1 Hybrid multi-mode microplate reader.

### Construction of recombinant plasmids

For the deletion of *divIVA* (DR_1369) and C-terminal domain (CTD from chromosome I in *D. radiodurans*, the recombinant plasmids pNOKUD4D and pNOKUND were constructed. For that, ∼1 kb upstream and downstream region from the coding region corresponding to full length, and CTD were PCR amplified using gene specific primers (Table S1) from the genome of *D. radiodurans*. PCR products were digested with required restriction enzymes and cloned in the pNOKOUT vector. The upstream and downstream fragment of *divIVA* were cloned at *Apa*I-*Kpn*I and *Bam*HI-*Xba*I sites in pNOKOUT (Kan^R^) to give pNOKUD4D. Similarly, 1.6 kb region covering NTD coding sequence of *divIVA* was replaced with upstream fragment in pNOKUD4D at *Apa*I-*Kpn*I sites to give pNOKUND. Similarly, pFtsZGFP expressing FtsZ-GFP on pVHSM plasmid under *P*_*spac*_ promoter; Spec^R^ (70 µg/ml) (Modi *et al*, 2014) was used for localization of FtsZ-GFP in wild type and Δ*divIVAC* mutant of *D. radiodurans*.

### Transformation in *D. radiodurans*

*D. radiodurans* was transformed with required plasmids using a modified protocol as described in (Udupa *et al*., 1994). In brief, *D. radiodurans* R1 was grown overnight in TGY broth and diluted to 0.3-0.4 OD_600_ with fresh TGY broth containing 30 mM CaCl_2_. This mixture was further incubated at 32°C for 1 hour. To this, 1–2 μg of circular or linearized plasmid was added per 1 ml of CaCl_2_ treated bacterial culture and cells were placed on ice for 45 min. The transformation mixture was incubated on an orbital shaker at 32°C for 30 min at 120 rpm. This transformation mixture was 10-fold diluted with TGY broth and grown for 15-18 h at 32°C in a shaker incubator at 180 rpm. Different dilutions of overnight grown cultures were plated on TYG agar plates supplemented with required antibiotic and plates were incubated at 32°C. The p11559 plasmid expressing spectinomycin resistance was used as a positive control and pSK+ conferring Amp^R^, was used as a negative control.

### Isolation of knockout mutants

For generating gene knockouts, the *D. radiodurans* cells transformed with *Xmn*I-linearized pNOKUD4D and pNOKUND plasmids were grown several generations in the presence of kanamycin (8μg/ml) as detailed in (Charaka and Misra, 2012). The transformants were scored on TGY agar plates supplemented with required antibiotic. Several independent transformants colonies were plated on TYG agar plate containing kanamycin. These cells were suspended in liquid broth and again plated on TGY agar supplemented kanamycin (8μg/ml). This cycle alternate streaking on TYG agar plate and sub-culturing in TYG broth was repeated at least 30-35 passages. Genomic DNA was prepared and homogenous replacement of target genes like *divIVA* and *divIVA*-C with the expressing cassette of *nptII* cassette was ascertained using diagnostic PCR. Since, *divIVA* replacement with *nptII* was indispensable, its homogenous replacement was not achieved. However, the homogenous replacement of CTD with expressing cassette of nptII could be achieved and these cells with a genotype *divIVA-C::nptII* were named as Δ*divIVAC* mutant.

### PCR screening of deletion mutation in *D. radiodurans*

Total genomic DNA of transformants was isolated as described earlier (Battista *et al*., 2001) and used as the template for diagnostic PCR. For identification and confirmation of disruption mutation, the gene-specific primers of antibiotic marker genes and target specific primers were used for PCR amplification in different combinations are given in Table S1. The diagnostic PCR was performed in two groups to verify the mutants. In one group, diagnostic PCR was performed using *ΔdivIVAC* mutant genomic DNA whereas in another group wild type genomic DNA was used. The different sets of flanking primers used for target genes and expressing cassette of kanamycin gene (*nptII*) were used as shown in Fig 2A and explained in figure legends. The PCR was performed using Phusion high-fidelity DNA polymerase and PCR products were analyzed on 1 % agarose gel after ethidium bromide staining.

### Immunoblotting of total cellular proteins

Immunoblotting of total proteins was carried out with specific antibodies as described earlier (Maurya *et al*., 2018). In brief, wild type and C-terminal domain mutant of *D. radiodurans* were grown overnight in TYG broth and TYG supplemented with kanamycin (8μg/ml) at 32°C. Next day, optical density at 600nm was measured online in the Synergy H1 Hybrid multi-mode microplate reader. Then, cells equivalent to OD_600_ 1.0 were taken, centrifuged and washed in 1X phosphate-buffered saline. Cell pellet was resuspended in equal volume of 10mM Tris-HCl pH8.0, 1mM EDTA and 2X Laemilli buffer. Cells were lysed at 95°C for 10 min and centrifuged. Equal volume of clear supernatant was run on 12% SDS-PAGE and blotted on PVDF membrane using BioRad Semi-dry blot machine. Blots were hybridised with polyclonal antibodies against DivIVA followed by anti-rabbit alkaline-phosphatase conjugated antibody and signals were developed using NBT-BCIP.

### Real time -PCR analysis

The quantitative Real time-PCR (qPCR) was performed for quantitative analysis of gene expression using LUNA universal one-step RT-qPCR kit and its protocol. In brief, total RNA was prepared from cells grown in TYG with antibiotic as required and qPCR was performed on QIAgen make Rotor-Gene Q real-time PCR cycler. Briefly, an equal amount of RNA (20 ng) was taken in 20 µl reaction mixture containing gene specific primers and other reagents as per the manufacturer’s instructions. RT-PCR was run at 55°C for 10 minutes for reverse transcription, 95°C for 1 minute for initial denaturation followed by 35 PCR cycles at 95°C for 10 seconds for denaturation, at 60°C for 30 seconds for extension and melt curve at 60–95°C with initial hold of 90 Sec at 60°C and 1°C raise for 1 second. Cq values were taken for calculation of the fold change in REST software.

### Fluorescence microscopy and image analysis

Fluorescence microscopy of different derivatives of *D. radiodurans* was carried out on Olympus Cell IX 83 microscope using standard procedures. In brief, the bacterial cultures were grown to stationary phase in TYG broth, fixed with 3 % paraformaldehyde for 10 min on ice and washed two times in phosphate buffered saline (pH 7.4). These cells were stained with Nile Red (0.5 µg/µl) and DAPI (0.5 µg/µl) for 10 min on ice and then washed three times with PBS. After washing, cells were resuspended in PBS, mounted on 1% agarose on glass-slides and samples were observed. Image analysis and cell size measurements were carried out using automated Cell Sens software incorporated in microscope. Similarly, wild type and mutant cells harbouring pFtsZGFP were grown overnight and 0.05-0.1 O.D. cells were taken for further growth in the presence of required antibiotics and induced with 20mM IPTG overnight. Next day, cells were processed for microscopy in the same manner described above. For expression of drDivIVA in *trans* in Δ*divIVAC* cells, pGroDivIVA expressing His-DivIVA under *groESL* promoter was transformed in these cells as described above (Udupa *et al*., 1994). Cells were grown in the presence of required antibiotic and processed for microscopy as mentioned above. For analysis, >200-300 cells stained with Nile red were taken from at least two separate microscopic fields of the images captured in two independent experiments and analysed for required attributes. Data obtained was subjected to Student’s t-test analysis using statistical programs of Graphpad Prism 5.0.

## Results

### Full length DivIVA deletion is indispensable in *Deinococcus radiodurans*

DivIVA is a membrane protein which has been shown to localise at cell poles in *B. subtilis*. The putative DivIVA in *D. radiodurans* (drDivIVA) is divided into N-terminal domain (NTD) between 1-165 amino acids and C-terminal domain (CTD) between 167-328 amino acids. Sequence comparison showed that its NTD is conserved and consists of lipid binding domain (Chaudhary *et al*., 2019) whereas CTD varies across Gram-positive bacteria. In order to understand *in vivo* role of drDivIVA, we attempted to make its null mutant by replacing with an expression cassette of *nptII* in *D. radiodurans* (Δ*divIVA*) as described in methods. We observed that the homogeneous replacement of *drdivIVA* with *nptII* has debilitated these cells when enrichment approached to homogeneity. These cells also showed the highly compromised growth under antibiotic selection. This observation was found to be true on both on TGY agar plate (Fig 1A) and in liquid medium (Fig 1B). Since, the CTD in drDivIVA is extra when compared with DivIVA of *Bacillus subtilis* and found to be variable across the Gram-positive bacteria, we got curious to probe CTD function(s) *in vivo*. For that, we made CTD deletion by replacement of *divIVAC* with *nptII* cassette through homologous recombination as described in methods. The perspective CTD mutant (Δ*divIVAC*) was viable under normal conditions on TGY agar plate in the presence of antibiotic (Fig 1C). The replacement of target sequence with nptII cassette was confirmed by diagnostic PCR using different set of primers as depicted in Fig. 1D). PCR product analysis confirmed the selective deletion of CTD domain of *divIVA* in chromosome I of *D. radiodurans* and integration of an expressing cassette of *nptII* in place of CTD coding sequences (Fig 1E). The Δ*divIVAC* cells were checked for growth in liquid medium, which grew slower than wild type (Fig 1F). These results together suggested that *divIVA* in *D. radiodurans* is an essential gene for its normal growth while NTD of DivIVA appear to be sufficient for its survival under normal conditions.

**Figure 1.**
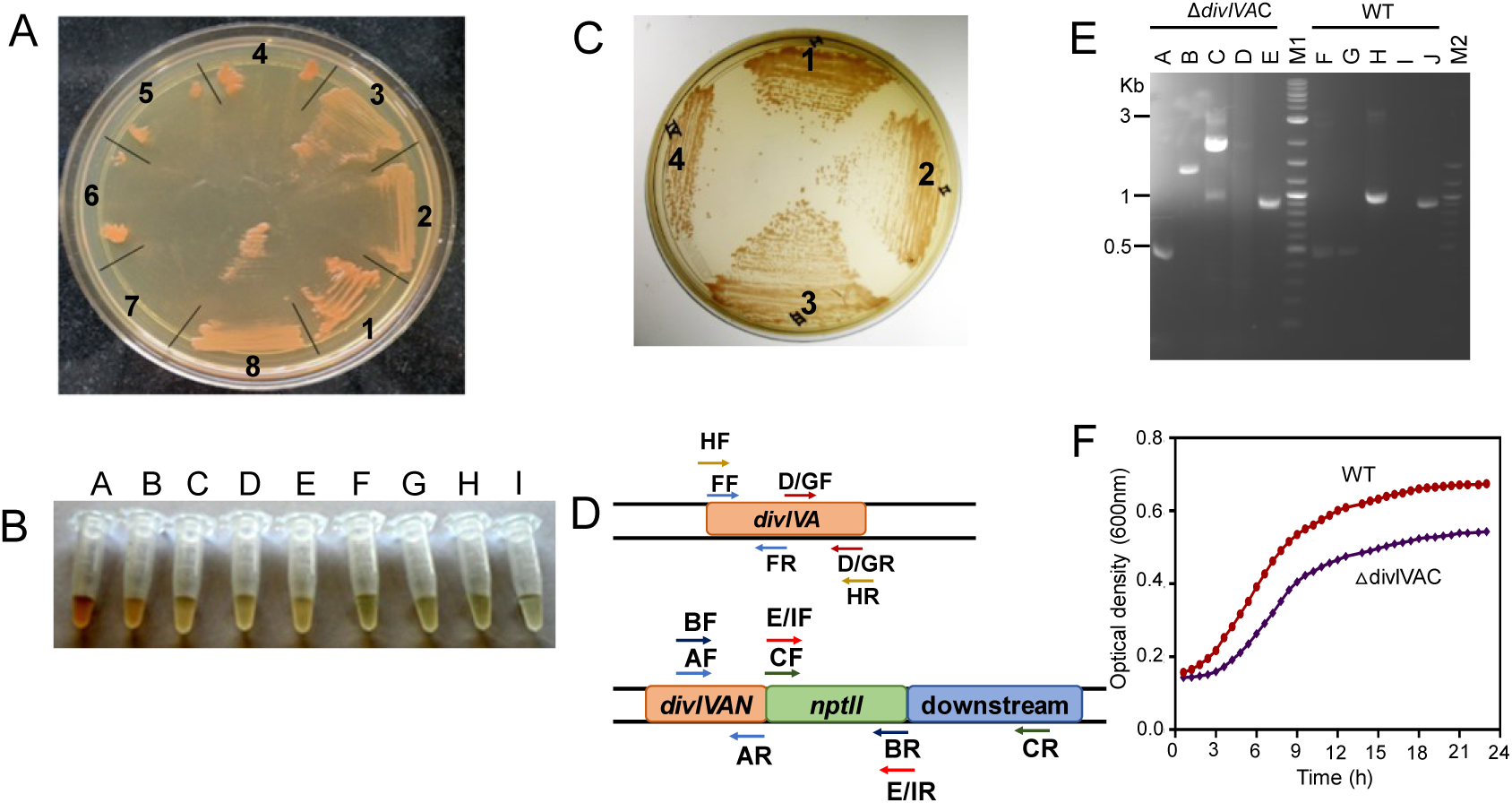
Growth characteristics of wild type *D. radiodurans* and its *divIVA* knockouts. *Deinococcus radiodurans* cells harbouring pNOKUD4D for creating *divIVA* knockout (A & B), and pNOKUND for creating CTD knockout (C) plasmids were maintained under antibiotics selection and expression of antibiotic resistance by way of *nptII* integration in place of target sequences into chromosome. For *divIVA* knockout, the transformants (1) were sub-cultured multiple times and kanamycin resistance was checked after 1^nd^ (2), 3^rd^ (3), 5^th^ (4), 7^th^ (5), 9^th^ (6) and 11^th^ (7) round of sub culturing and their growth was compared with *ΔparA* (8) mutant as positive control (A). Similarly, transformants grown on plate were also checked for their survival in liquid culture (B). Tubes C-I represents the growth conditions of these transformants sub-cultured for 9 rounds whereas tube A and B show growth of wild type and *ΔparA* respectively. Similarly, CTD mutant (Δ*divIVAC*) cells were also grown on TYG plate supplemented with required antibiotic from different 2^nd^ (1), 4^th^ (2), 6^th^ (3) and 10^th^ (4) rounds of its sub culturing showing normal growth (C). The perspective Δ*divIVAC* mutant after 10^th^ rounds of sub-culture was diagnosed for homogenous replacement of CTD with *nptII* cassette by PCR amplification using genomic DNA from mutant (Δ*divIVAC*) and wild type (WT) and primers as schematically depicted (E) and detailed in Table S1. The PCR products obtained from AF and AR (A), BF and BR (B), CF and CR (C), D/GF and D/GR (D & G), E/IF and E/IR (E & I), FF and FR (F), HF and HR (H) sets of primers were analysed in agarose gel (E). The normal growth of confirmed Δ*divIVAC* mutant was compared with wild type (F).

### CTD has an important role in drDivIVA expression

Immunoblotting of total proteins from the wild type and Δ*divIVAC* cells was carried out using polyclonal antibodies against drDivIVA, to check the correctness of CTD deletion in the chromosome and the expression of full length and NTD in these cells was done by immunoblotting using drDivIVA antibodies. The results confirmed the expression of full length drDivIVA in the wild type and only NTD in Δ*divIVAC* cells (Fig 2A). Interestingly, we noticed that the level of NTD was ∼10 fold more in Δ*divIVAC* cells than full-length drDivIVA in wild type. Since, upstream region of *divIVA* in both wild type and Δ*divIVAC* mutant is same and difference lies only in their CTD, the possibility of CTD affecting either transcription or stability of full-length protein could be hypothesized. To answer it, the transcript level of *divIVAN* was measured in the both mutant and wild-type cells and compared with the levels of *minC* and *minD* expression in similar backgrounds. Results showed that the transcript levels of *divIVAN* were ∼14 times higher in Δ*divIVAC* cells than wild type cells (Fig. 2B). Although, this observation is highly intriguing, a strong possibility of CTD role in autoregulation of drDivIVA level in *D. radiodurans* could be articulated. These results suggested that CTD plays an important role in the regulation of DivIVA expression in this bacterium and regulated expression of DivIVA is first time shown in any bacteria.

**Figure 2:**
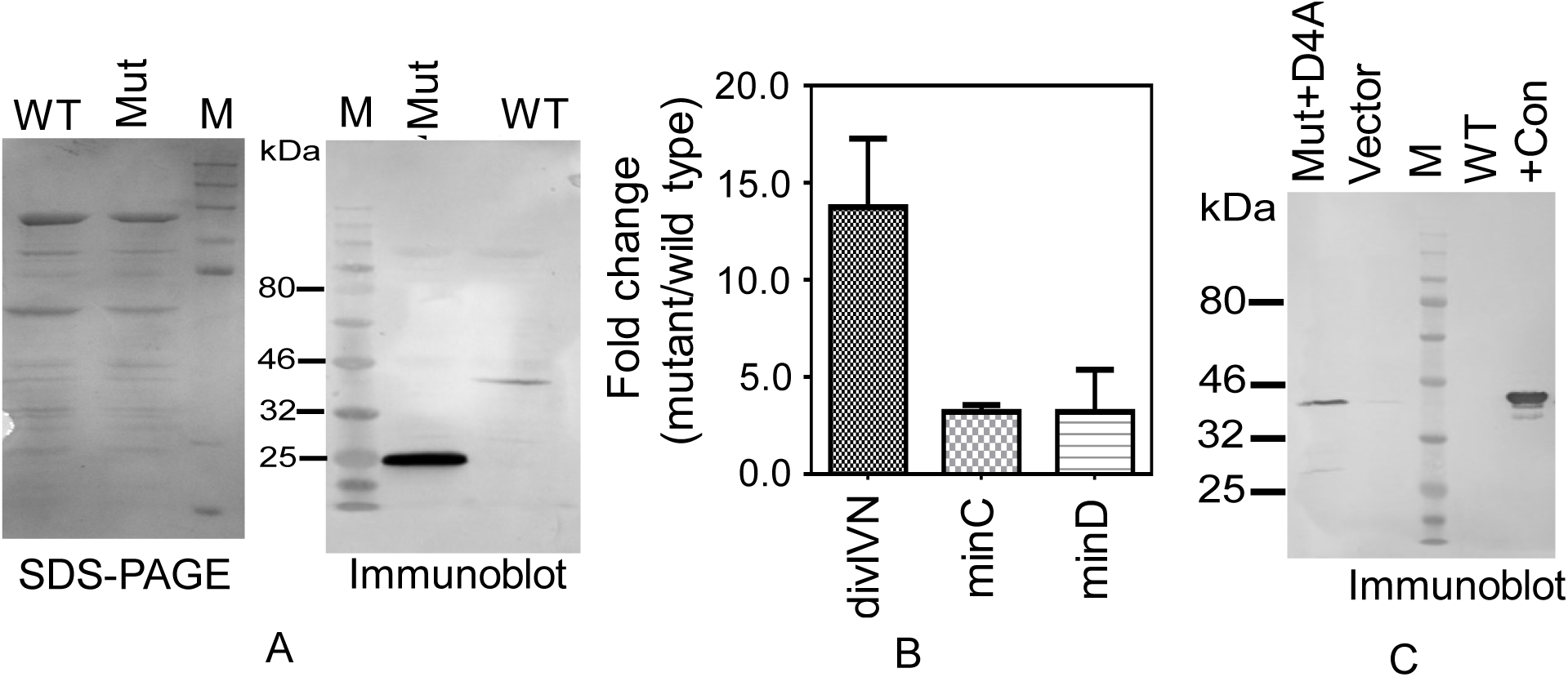
Confirmation of CTD deletion and expression of DivIVA in *D. radiodurans*. The *D. radiodurans* R1 (WT) and its Δ*divIVAC* mutant (Mut) were grown and total proteins were separated on SDS-PAGE (SDS-PAGE). These proteins were immunoblotted with antibodies against DivIVA protein (immunoblot) (A). Similarly, total RNA was prepared from both WT and Δ*divIVAC* mutant (Mut) and the levels of NTD (divIVN) of DivIVA, *minC* (minC) and *minD* (minD) transcript were estimated using RT-qPCR. Fold change in the transcript of these genes was estimated using REST software (B). The Δ*divIVAC* mutant cells were transformed with wild type DivIVA expressing plasmid (Mut+D4A), empty vector (Vector) and levels of DivIVA fused with his tag was estimated using anti-his antibodies and compared with wild type (WT) and positive control having his tagged DivIVA (+Con) (C). Sizes of protein bands were compared with molecular weight marker (M).

### CTD deletion mutant produces bent septum in tetrad colonized cells

Since, DivIVA is member of Min system in Gram-positive bacteria and its role in septum formation in known in other bacteria, we examined Δ*divIVAC* cells for morphological change if any, under fluorescence microscope. The Δ*divIVAC* mutant showed tetrad colonization where each compartment contained DNA similar to wild type cells. However, when we measured the cell surface area using Olympus Cell Sens software, it was found that mutant cells with their average size of 4.8 ± 0.18 µm^2^ were significantly (P < 0.0001) smaller than wild type cells having an average size of 6.5 ± 0.36 µm^2^ (Figure 3A). Interestingly membrane staining of Δ*divIVAC* mutant with Nile Red showed that ∼70 % population has bent septum as compared to less than 5% wild type with similar phenotype. Similarly, ∼30 % of mutant population showed normal septum which was observed in ∼95 % of wild type cells (Figure 3B). Curiously, we analysed the angle of septum bend from its normal division plane with respect to previous division plane. In mutant cells new septum is formed at ∼70° angle from the plane of previous septum, which was significantly different with P < 0.0001 from the ∼90°angle as observed in wild type (Fig 3C). Notably, we did not detect mini cells like phenotype as observed in rod shaped bacteria, and all the cells in the population contain DNA. These results might suggest CTD role in the regulation of DivIVA mediated perpendicular growth of new septum in this bacterium.

**Figure 3:**
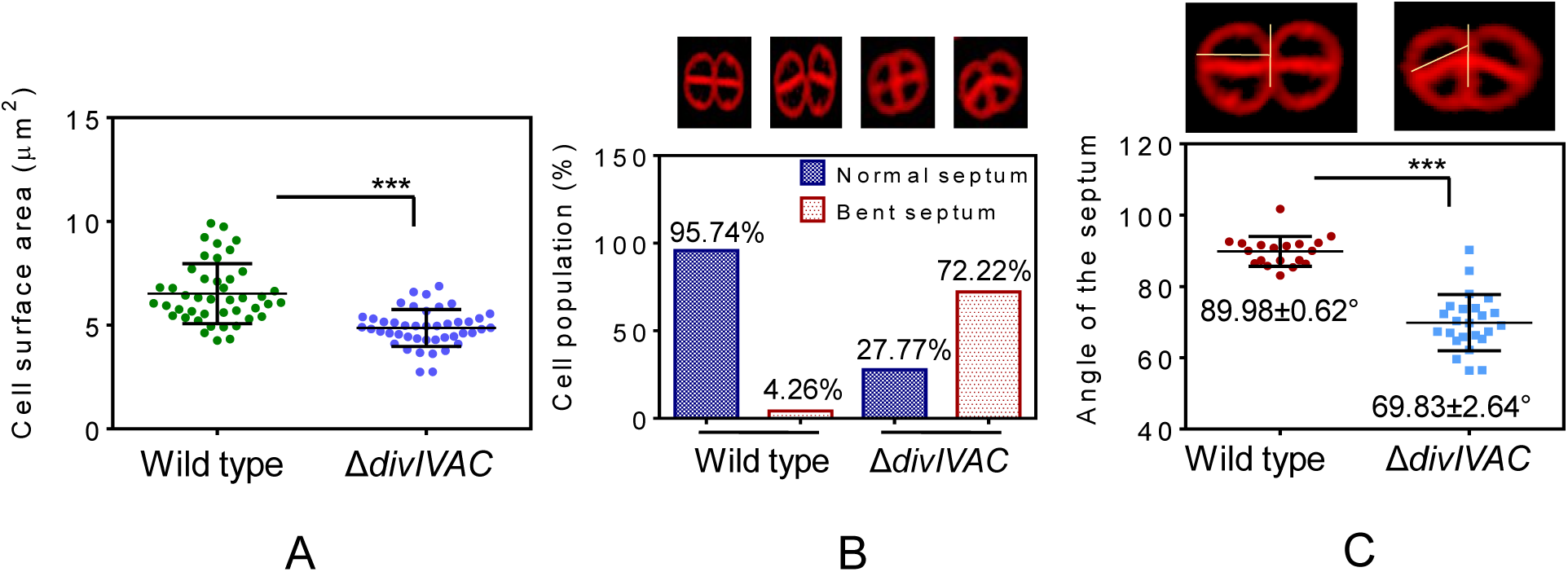
Effect of CTD deletion on cell size and septum characteristics in *Deinococcus radiodurans*. Both wild type and CTD mutant (Δ*divIVAC*) were grown overnight, stained with DAPI and Nile red, and observed under fluorescence microscope. Nearly 250 cells were measured for cell surface area using Olympus Cell Sens software and analysed using Student’s t-test (A). Similarly, these cells were also observed for the physical appearance of the septum like straight (normal septum) or tilted (bent septum) as shown in images of upper panel (B) and also the angle of the septum with respect to previous septum as marked in the images given in upper panel (C). Results were analysed employing Student’s t-test. *** denotes p values <0.0001.

### FtsZ mis-localized in Δ*divIVAC* mutants

The morphological abnormalities in size and septum placement in Δ*divIVAC* mutant were further studied by monitoring FtsZ localization in these cells. For that, FtsZ-GFP was expressed in both wild type and Δ*divIVAC* cells and FtsZ localization was monitored using fluorescence microscopy. We observed that 2 juxtaposed foci of FtsZ-GFP flanking to middle foci were not in straight line rather at an angle in Δ*divIVAC* while these were in straight line in wild type cells (Fig 4, WT). The position of FtsZ-GFP was further analysed by Line Scan Analysis (LSA). We observed a normal pattern of FtsZ localization at mid cell position in wild type cells while FtsZ-GFP localization in mutant was grossly distorted and foci were away from mid cell position (Fig 4, Δ*divIVAC*). Nearly 250 cells of both mutant and wild type were analysed for cell length from one end to other longitudinally, and width across septum by fixing FtsZ foci position as an origin point, and the values were plotted as an average cell length on X-axis. Results showed that FtsZ foci are on average placed closed to axis in wild type cells (Fig 5A) while these were scattered to one side away from axis in Δ*divIVAC* mutant (Fig 5B). We further checked the effect of CTD deletion on FtsZ functional integrity like localization and polymerization. When we did Z-stacking and 3D images were constructed, both wild type and mutant cells showed FtsZ localization at juxtaposed position in membrane *albeit* they were positioned away from the perpendicular axis of septum and produced seemingly functional Z ring structure (Fig. 5 C). These results suggested that although CTD deletion has not affected FtsZ localization and polymerization *per se* but has affected the proper positioning of FtsZ-ring and its geometrical symmetry in mutant cells by yet unknown mechanism.

**Figure 4:**
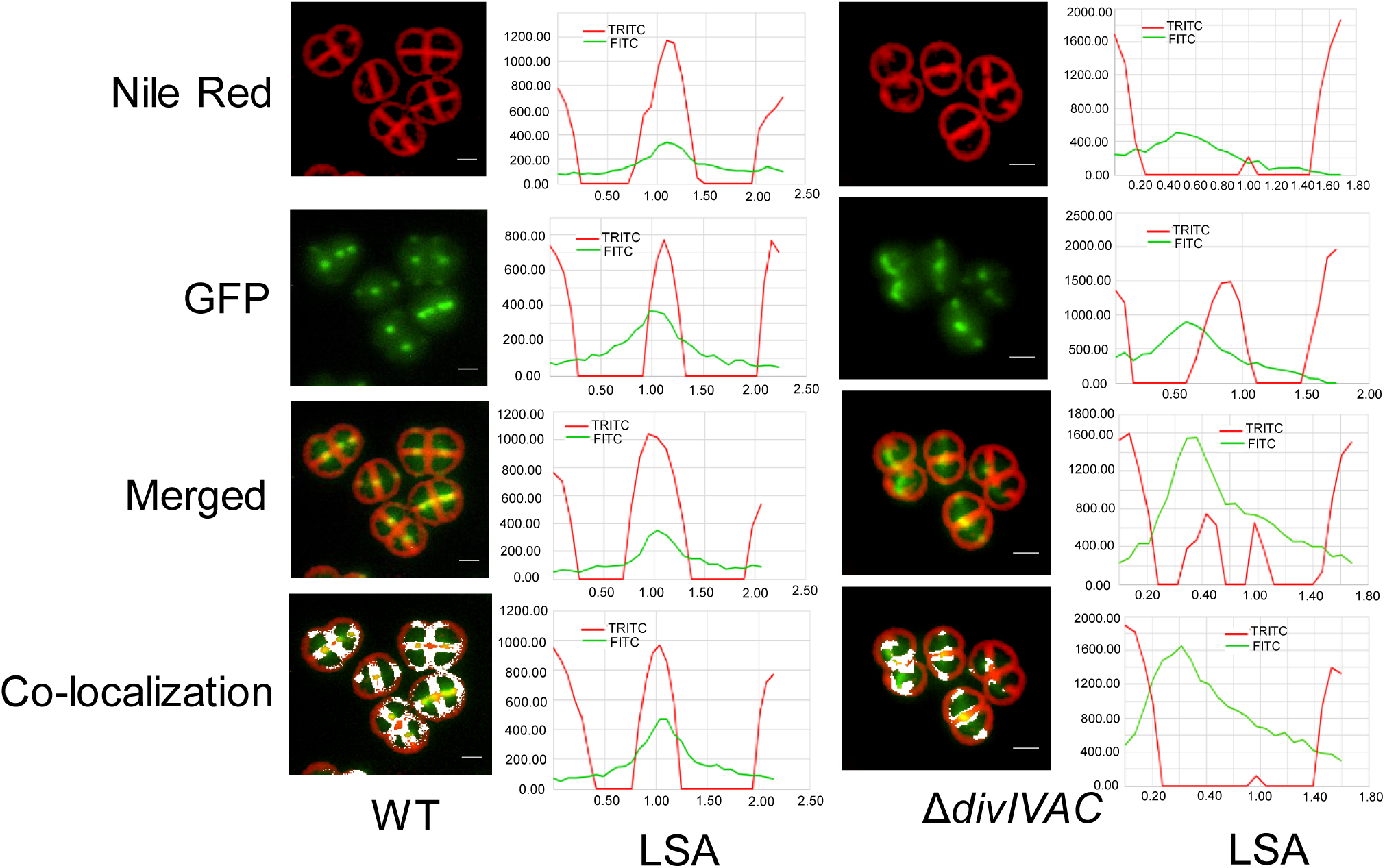
Effect of CTD deletion on FtsZ localization in *Deinococcus radiodurans*. The wild type (WT) and CTD mutant (*ΔdivIVAC*) cells expressing FtsZ-GFP fusion under inducible *spac* promoter were stained with Nile red and imaged under TRITC (Nile Red) and FITC (GFP) channels. These imaged were merged (Merged) and the colocalization patterns of septum with FtsZ-GFP was analysed (Co-localization). The data shown here is selectively for FtsZ-ring positioning in diad cells. Line scan analysis was done in a diad cells from one point of the cell to the other end using Olympus CellSens software, as detailed in methods and the position of the Z-ring in respective samples was examined and shown beside respective images (LSA).

**Figure 5:**
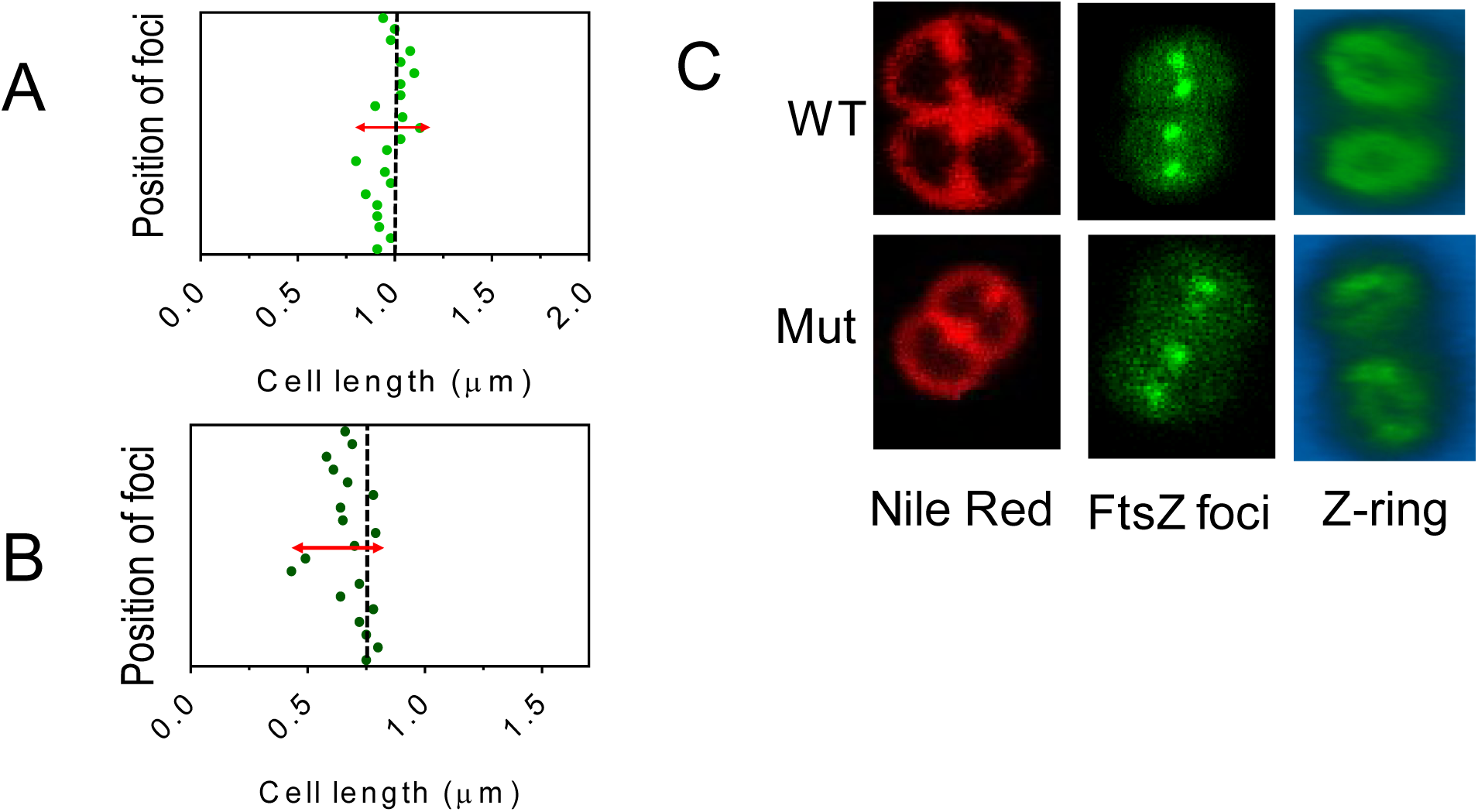
Comparative analysis of FtsZ localization and polymerization in wild type and CTD mutant of *Deinococcus radiodurans*. The cell length of *D. radiodurans* (A) and its CTD mutant (B) from FtsZ-GFP foci was measured and plotted as a function of cell length measured from one end to another end in nearly 250 cells. The red arrows indicate the FtsZ position deviation from central axis. The Z stacking of cells with FtsZ-GFP foci was done from its origin point for wild type (WT) and CTD mutant (Mut) and 3-D images of Z ring was constructed using image J software (C).

### Complementation of Δ*divIVAC* phenotype by drDivIVA *in trans*

To confirm that the observed phenotype was caused by the deletion of CTD in Δ*divIVAC* cells, we expressed drDivIVA in Δ*divIVAC* cells on pGroDivIVA under a constitutive *groESL* promoter (Table S1) and monitored under fluorescence microscope mainly for septum architecture. We observed the nearly complete restoration of wild type septum growth that was perpendicular to the plane of previous septum, and cell size of mutant cells recovered to near wild type (Fig. 6). However, a large number of mutant cells expressing wild type protein showed defect of delayed resolution of tetrad from octet colonies. Since, the levels of drDivIVA was significantly high in complemented population (Fig. 2C), so this phenotype could be implicated to higher level of wild type protein in these cells and that supports the autoregulation of drDivIVA as suggested to its CTD contribution, for the normal growth of this bacterium.

**Figure 6:**
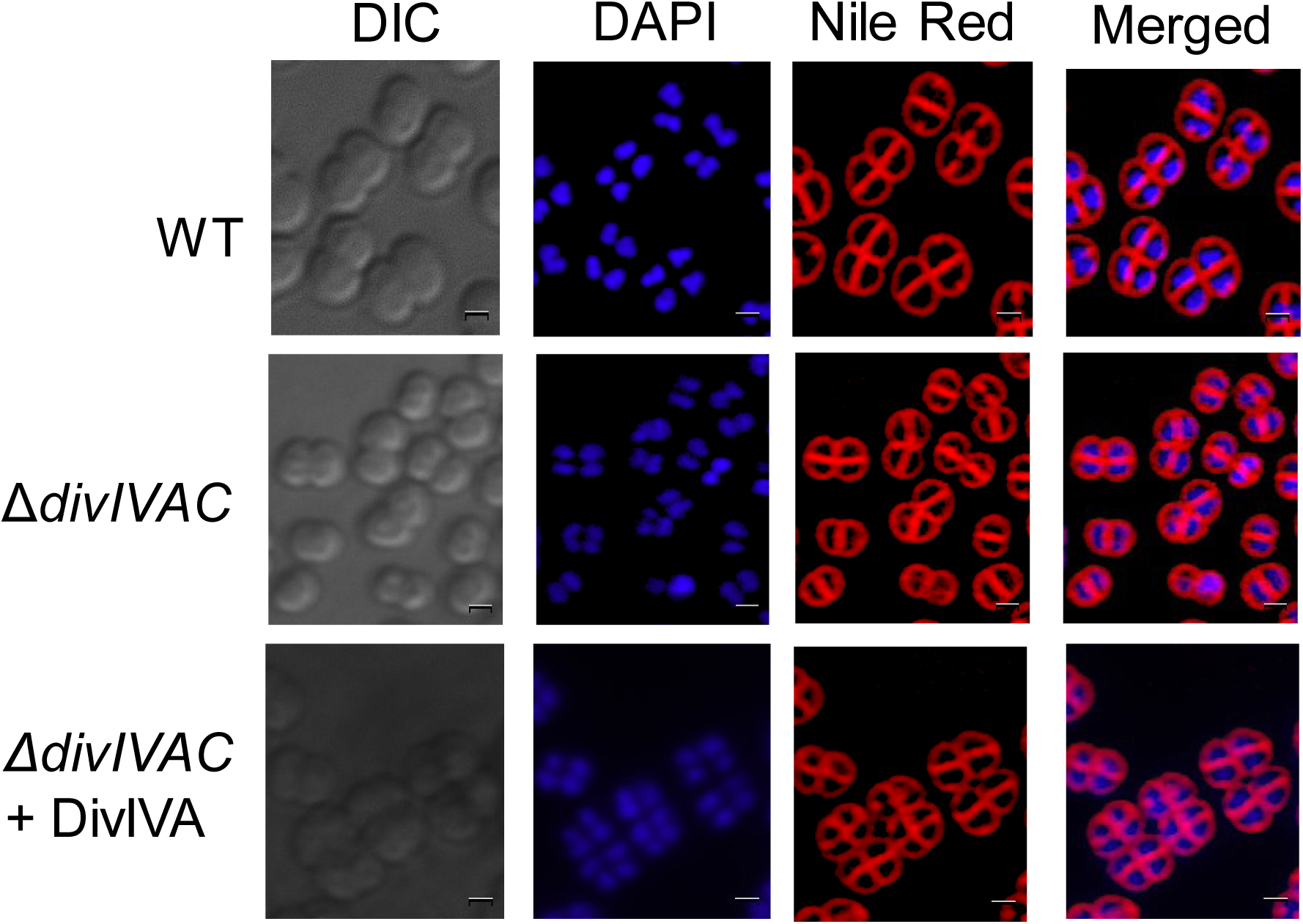
Functional complementation of DivIVA in CTD mutant of *Deinococcus radiodurans*. The wild type (WT), CTD mutant (Δ*divIVAC*) CTD mutant expressing DivIVA on plasmid (*ΔdivIVAC +* DivIVA) cells were stained with DAPI and Nile Red and imaged microscopically. These images were examined for the position of new septum with respect to previous septum. Data shown is a representative of the reproducible experiments.

### DivIVA localizes in membrane and perpendicular to alternate plane of cell division

The wild type cells expressing DivIVA-RFP under *groESL* promoter were stained with fluorescent labelled vancomycin-BODIPY dye, DAPI and observed under fluorescence microscope. The vancomycin stained septum with green fluorescence distinguishes the septum from red fluorescent foci of DivIVA-RFP. We observed DivIVA localization in the membrane (Fig 7A). Careful examination of these images showed that each DivIVA foci was aligned with either old septum or newly growing septum in the cell. The positions of these foci were at a regular space and almost perpendicular to each other. Interestingly, the formation of new DivIVA-RFP foci in membrane was oriented to both perpendicular and parallel to the direction of duplicated genome segregation in alternative planes of cell division. These results suggest the role of DivIVA in polarity determination, which seems to be in tandem with genome segregation.

**Figure 7:**
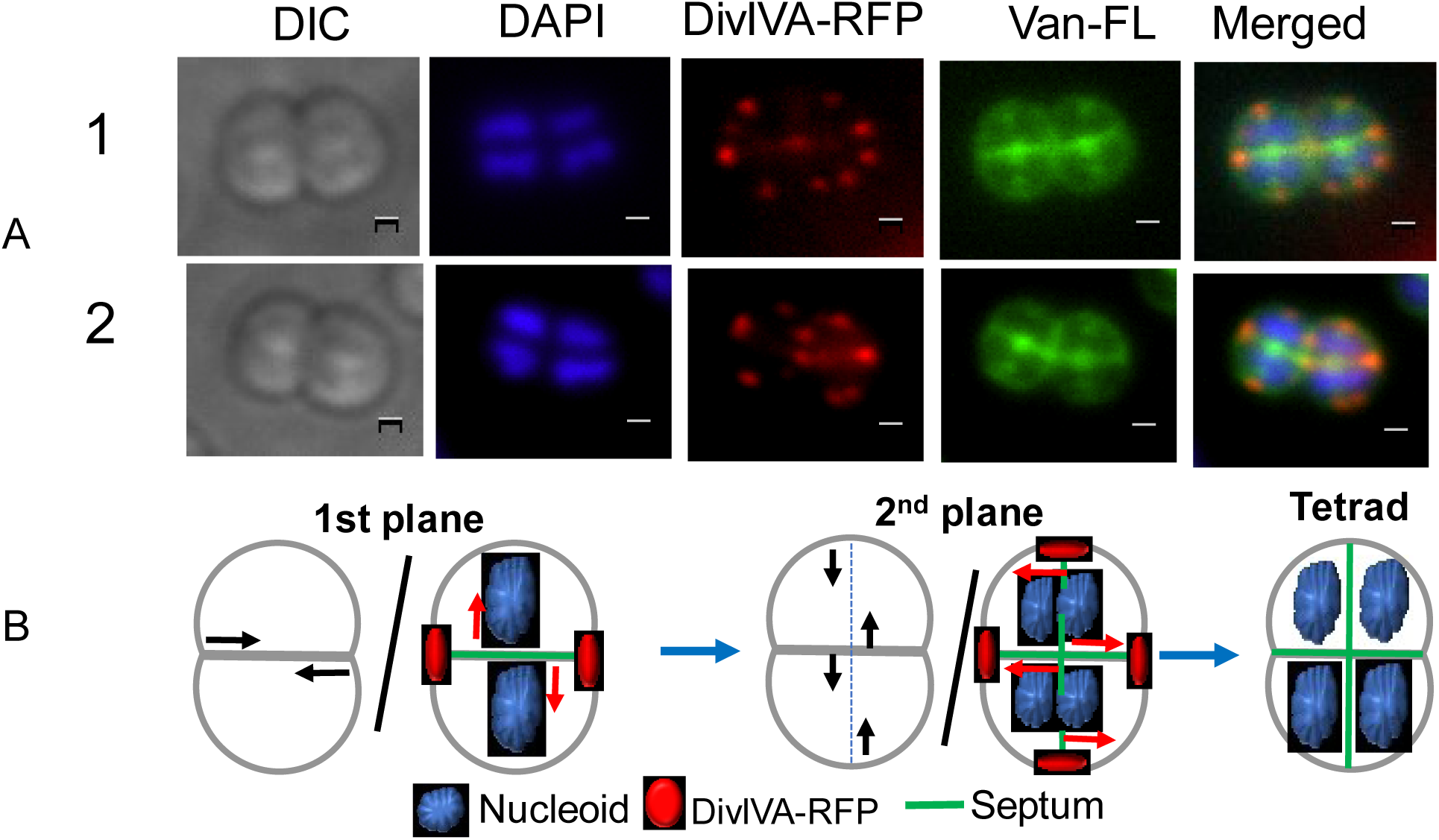
Localization of DivIVA-RFP in dividing population of *Deinococcus radiodurans*. The wild type cells expressing DivIVA-RFP under a constitutive *groESL* promoter were stained with DAPI and vancomycin-fluorescent-BIODPY and examined under fluorescence microscope for nucleoid (DAPI), septum (Van-FL) and DivIVA (DivIVA-RFP) localization. Data shown are representatives of two independent reproducible experiments (1 and 2) (A). For better clarity of the data obtained from majority of the cells, the typical pattern of DivIVA-RFP localization in cells moving from first cell division plane (1^st^ plane) and next round of cell division (2^nd^ plane) before producing tetrad cells are depicted schematically (B).

## Discussion

*Deinococcus radiodurans* a round shaped tetrad colonized radioresistant bacterium divides in alternate planes and the new plane is perpendicular to the previous plane of cell division (Murray *et al*., 1983). Unlike rod shaped bacteria, molecular basis that determines the cell position for FtsZ localization for septum initiation, which would be perpendicular to previous plane is very little understood in cocci and has not been studied in *D. radiodurans* yet. The roles of Min system in general and NOC system in particular to DNA metabolism have been implicated as the major regulator of spatial and temporal event associated with the initiation of FtsZ polymerization at the mid cell position in rod shaped bacteria. The *D. radiodurans* genome encodes MinC, MinD and both MinE though aberrant, and DivIVA (White *et al*., 1999). DivIVA belongs to Min protein family where they identify and determine the poles in the cells and allow FtsZ ring to form in mid-cell position in rod shaped Gram-positive bacteria (Thomaides *et al*., 2001). In *E. coli*, mid cell position is regulated by oscillation of MinC proteins and there are many key factors that regulate pole-to-pole oscillations of MinC protein. These are; (i) the highly negative membrane curvature at the poles (Renner & Weibel, 2011), (ii) the longest axis of confinement (Schweizer *et al*., 2012), (iii) the longest possible distance for the diffusion of MinD (Corbin *et al*., 2002), and (iv) the symmetry axes and scale of the cell shape (Wu *et al*., 2015). Also, in *E. coli*, MinC oscillates from one pole to another being dependent on oscillatory patterns of MinD and MinE (Hu *et al*., 1999). It has been shown that DivIVA in general detects negative curvature at the cell poles, which is marked by cardiolipin domains in *B. subtilis* and thus localizes at the poles. NTD in drDivIVA has a lipid binding motif and is structurally supported by an extra CTD when compared with *B. subtilis* DivIVA. We showed that CTD contributes in the stabilization of a functional structure to drDivIVA (Chaudhary *et al*., 2019). In addition, the *D. radiodurans* cells being coccus shape elongates along the axis and attains to elliptical shape without any major change along central septum. In such a scenario, if DivIVA senses the geometry of the cells, marking negative curvature at the poles for MinC to oscillate in parallel to the central septum in this bacterium, cannot be ruled out. Additionally, it has been demonstrated that DivIVA interact with proteins involved in genome maintenance and those regulates cell division (Maurya *et al*., 2016, Chaudhary *et al*., 2019). These anticipated morphological changes in *D. radiodurans* during cell division, and drDivIVA interaction with macromolecular complexes associated with cell division and genome segregation together suggest the possible mechanisms by which drDivIVA could help in cell polarity determination in this bacterium.

Here, we present results that began providing evidences on the roles of drDivIVA in physiology of *D. radiodurans*. When we attempted to create deletion of full length *drDivIVA*, it was understood that DivIVA is an essential protein in *D. radiodurans* (Fig 1). Essentiality of divIVA for bacterial survival had been different in different bacteria. For instance, the DivIVA was indispensable in *Enterococcus faecalis* (Ramirez-Arcos *et al*., 2005) whereas it could be deleted in *Staphylococcus aureus* without loss of viability (Pinho & Errington, 2004). Since, CTD of DivIVA is variable across the DivIVA of different bacteria and NTD is conserved, we created CTD deletion in the genome of this bacterium. We observed a clear phenotype in cell lacking CTD that include the smaller cell size and bent septum when compared with wild type. To understand the reason of bent septum, we checked FtsZ localization in both wild type and CTD mutant cells. We noticed that CTD mutant fails to guide the proper localization of FtsZ required for septum growth. This resulted FtsZ-GFP juxtaposed foci formation at an angle, which were in straight line in wild type cell. The possible reasons of angular FtsZ ring formation in CTD mutant instead of the perpendicular type observed in wild type is not known yet but might have indicated DivIVA role in determining the position of septum growth. Earlier, we know that drDivIVA interacts with MinC through its middle domain which apparently folds correctly only in full length protein but not when either NTD or CTD is missing from the protein. The oscillation of Min system along the long axis of the ellipsoidal shaped cell and in parallel to the central septum if determines the precise placement for septum growth during cell division cannot be ruled out. In rod shaped bacteria, the longitudinal oscillation of MinC between the poles, helps FtsZ to localize in the centre and forms FtsZ ring perpendicular to the oscillatory path of MinC. In *D. radiodurans*, drDivIVA binds to membrane through its N-terminal lipid binding motifs and MinC by intracellular middle domain, which is part of CTD. These two facts together might allow us to propose a model that the longitudinal oscillation of MinC is perhaps regulated by MinC interaction with drDivIVA, which controls the precision in longitudinal oscillation of MinC and hence proper localization of FtsZ and normal septum growth. In the absence of CTD, the membrane anchored NTD does not interact with MinC and hence its oscillation might lose the precision in longitudinal dynamics and if that happens, the FtsZ localization will also get distorted, which could affect the architect of new septum (Fig 8). Since, drDivIVA also interacts with genome segregation proteins through its NTD, the possibility of drDivIVA interaction with genome segregation proteins if affecting perpendicularity of septum growth cannot be ruled out. The wild type cells expressing DivIVA-RFP under constitutive promoter showed DivIVA localization in the membrane aligned with old septum as well as in membrane aligned for next division plane of cell division (Fig. 7). Careful examination of vancomycin and DAPI stained cells showed that DivIVA-RFP foci in second plane of division was perpendicular to direction of DNA segregation. DivIVA interaction with proteins other than Min system has also been reported and the impact of such interactions on bacterial cell physiology and growth has been suggested (Ginda *et al*., 2013). Therefore, a possible role of DivIVA interaction with genome segregation proteins could be to mark the new plane for septum formation, which would be parallel to earlier direction of genome segregation but perpendicular to old septum.

**Figure 8:**
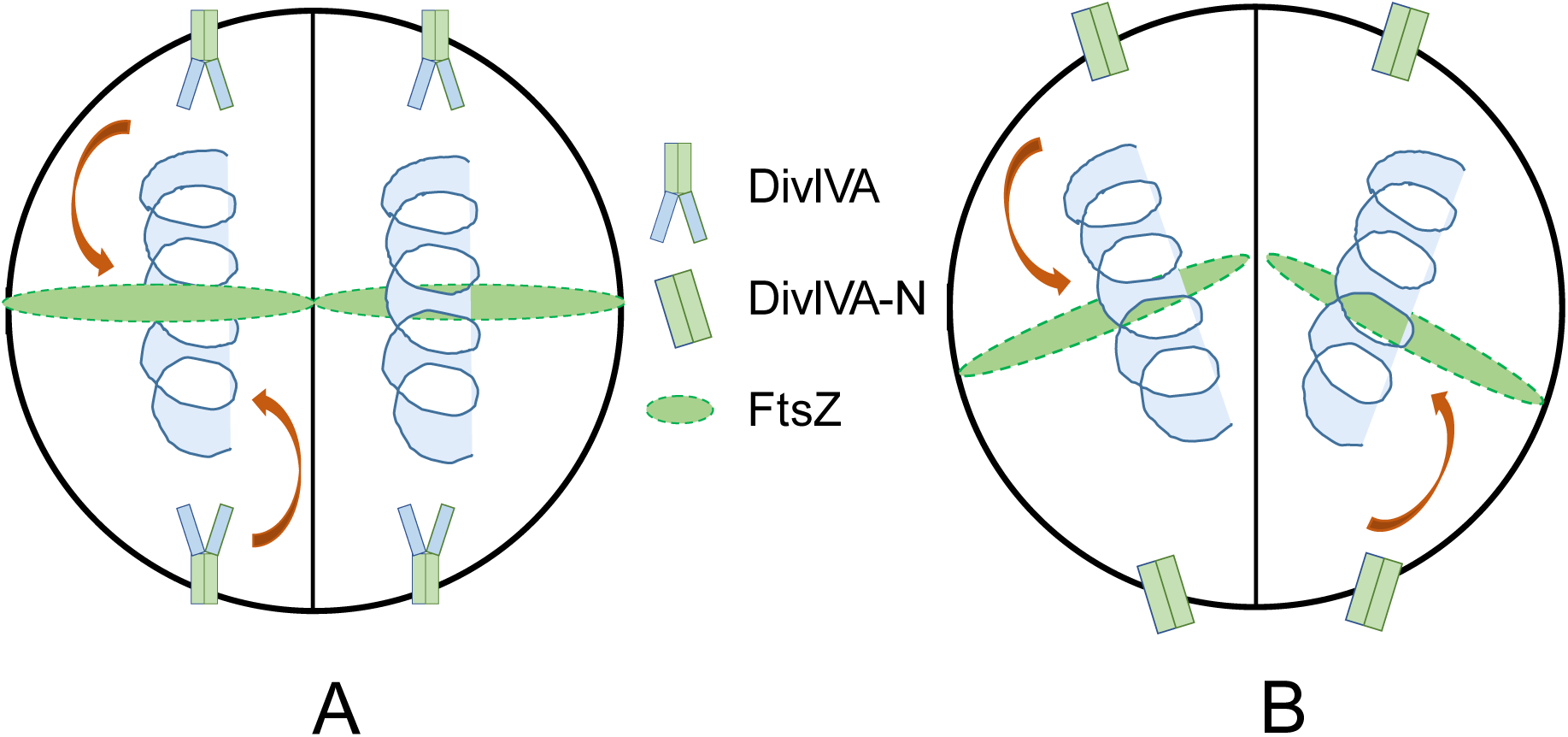
A schematic model explaining MinC oscillation in wild type and CTD mutant of *Deinococcus radiodurans*. Model shows membrane localization of full length DivIVA (A) in wild type and its NTD variant (DivIVA-N) (B) in CTD mutant and possible differences in the interaction with MinC that would arise due to absence of CTD in DivIVA-N protein. This model may explain that how MinC oscillation through the longitudinal axis in wild type or away from it, in the absence of CTD can be a deterministic factor for perpendicular (A) and angular (B) growth of FtsZ-ring mediated new septum. Symbols representing DivIVA, DivIVA-N and FtsZ are shown for better understanding.

In summary, we report that *D. radiodurans* failed to tolerate genomic deletion of full length divIVA suggesting its essentiality in the normal growth of this bacterium. Interestingly, the selective deletion of CTD encoding sequence from the genome was possible and the mutant cells showed scorable phenotypes like smaller size cells, bend septum and delayed growth when compared with wild type cells. Although, molecular basis of essentiality of drDivIVA and how NTD in the absence of CTD is accounting for bent septum phenotype and reduction in cell surface area are not clear and can be addressed independently, the available results provide a clear evidence on indispensability of drDivIVA in *D. radiodurans* and CTD role in determining the position of new septum perpendicular to previous one, most likely through its interaction with genome segregation proteins.

## Acknowledgements

We thank Mrs. K.V. Boby TIFR for her help in confocal microscopy. Reema Chaudhary is grateful to Department of Atomic Energy (DAE), Government of India for the research fellowships.

