## Supplementary material for "DivIVA is essential in *Deinococcus radiodurans* and its C terminal domain regulates new septum orientation during cell division": Table S1 and TableS2

Supplementary Table S1 and S2

**Table S1. List of Primers used in this study**

| Primer Name | Oligonucleotide Sequences | Purpose | Name used for diagnostic PCR |
|---|---|---|---|
| D4AUPFw | 5’ CG GGGCCC ACGGGAATGCTCACGGCCGCC 3’ | pNOKUD4D, pNOKUND | - |
| D4AUPRw | 5’ CG GATATC CGGATTATTGTTATTGGGCGA3’ | pNOKUD4D | - |
| D4ADNFw | 5’ CG GGATCC AAACGGCCAGGCCGTCGT 3’ | pNOKUD4D, pNOKUND | - |
| D4ADNRw | 5’ GC TCTAGA TCTCGGCCTGAGCACTGGC 3’ | pNOKUD4D, pNOKUND | CR |
| D4PETF | 5’ CG GGGCCC ATGAGCTCGCCAATAAC 3’ | diagnostic PCR, qPCR | AF, BF, FF, HF |
| D4NPETR | 5’ CG GAATTC TTACGAGCCGCTCACCCT 3’ | pNOKUND, diagnostic PCR, qPCR | AR, FR |
| D4CPETF | 5’ CG GGATCC ATGGACCTCGAGCGGCAGTT 3’ | diagnostic PCR | D/GF |
| D4PETR | 5’ CG GAATTC TTATTTCTCGTCGTCCAGC 3’ | diagnostic PCR | D/GR, HR |
| nptIIFw | 5' GCACGGTGGCCGAGTGG 3' | diagnostic PCR | CF, E/IF |
| nptIIRw | 5' GTCAGCGTAATGCTCTG3' | diagnostic PCR | BR, E/IR |
| MC F | 5' CAATGCGAATCTGAGCACTGGCGA T 3' | qPCR | - |
| MC R | 5' CGGTGGAAATTCAGGGCGACACTCA 3' | qPCR | - |
| MD F | 5' ACCATGTCGTCGATGGAGAGCATGTT 3' | qPCR | - |
| MD R | 5' AAGTGCCGCATGAATCAGGCACTG A 3' | qPCR | - |
| pETHisFw | 5’ AAAAGTACTGGGCCCATGGGCAGCAGCCAT 3’ | pGroDivIVA | - |
| pETHisRw | 5’ CGCTTAAGTCTAGATATCTCAGTGGTGGTG 3’ | pGroDivIVA | - |

### Table S2: List of bacterial strains and plasmids used in this study

| Bacterial strains | Genotype | Source |
|---|---|---|
| *Deinococcus radiodurans* R1 | Wild type strain ATCC13939 | Lab Stock |
| *E. coli* NovaBlue | *end*A1 *hsd*R17$(r_{K12}^{-}\ m_{K12}^{+})$ *sup*E44 *thi-1* *rec*A1 *gyr*A96 *rel*A1 *lac*F''*[pro*$A^{+}B^{+}$ *lac*$I^{q}$ *Z*ΔM15*::*T*n*10 ] ($Tet^{R}$) | NEB Inc., |

**Plasmids**

| Sr No. | Plasmids | Characteristics | Sources | MW of Protein (~kDa) |
|---|---|---|---|---|
| 1 | pNOKOUT | $pBSK^{+}$ ($Amp^{R}$) containing *nptII* casette (937 bp) at SmaI site, ~ 4.0 kb | Khairnar *et al.,* 2008 | - |
| 2 | pNOKUD4D | pNOKOUT containing upstream (~ 1.0 kb) and downstream (~ 4.0) kb sequences of dr*divIVA*, $Amp^{R}$/ $Kan^{R}$ | This study | - |
| 3 | pNOKUND | pNOKOUT containing upstream + *divIVA-N* (~ 1.6 kb) and downstream (~ 1.0) kb sequences of dr*divIVA*, $Amp^{R}$/ $Kan^{R}$ | This study | - |
| 4 | pRadgro | pRAD1 containing *groESL* promoter (261 bp) at BglII-XbaI site, ~ 6.5 kb, $Amp^{R}$ | Misra *et al.,* 2006 | - |
| 5 | pGroDivIVA | pRadgro containing *his-divIVA* at ApaI-XbaI site, ~ 7.7 kb, $Amp^{R}$ | This study | ~40 kDa |
| 6 | pFtsZGFP | pVHSM containing ftsZ-gfp at SacI-AflII site, ~ 11.7 kb, $Spec^{R}$ | Modi *et al.,* 2015 | ~40 kDa |
